# *In vitro* and *in silico* analysis proving DPP4 inhibition and diabetes associated gene network modulation by a polyherbal formulation – *Nisakathakadi Kashaya*

**DOI:** 10.1101/2022.07.15.500175

**Authors:** Anjana Thottappillil, Sthitaprajna Sahoo, Abhijnan Chakraborty, Sania Kouser, R. Vidhya Ravi, Soumya Garawadmath, Pranav Girish Banvi, Subrahmanya Kumar Kukkupuni, Suma Mohan S, Chethala N Vishnuprasad

## Abstract

Frontiers of disease biology started recognizing the importance of systems and network medicine approach for managing chronic disease like diabetes. Dipeptidyl-peptidase IV (DPP4) inhibitors are one such class of anti-diabetic drugs recognized for their systemic biological actions. Polyherbal preparations like *Ayurveda* formulations are ideal for identifying novel DPP4 inhibitors having greater efficacy and safety profile. Additionally, expanding the research on the multitargeted mode of action of these polyherbal formulations can render novel insights into the complex biology of disease manifestations. The current study aims at identifying DPP4 inhibitory potential of a clinically established Ayurveda anti-diabetic formulation *Nisakathakadi Kashaya* (NK) using *in vitro* and *in silico* methods as well as the modulation of diabetes associated gene network by NK. a. Standard enzyme inhibition assay was used to study the DPP4 inhibitory potential of NK, followed by bioinformatics and computational biology tools for identifying the potential bioactives and their molecular interactions involved in DPP4 inhibition. STITCH, CHEMBL and BindingDB databases were used for target mapping and depicting the multi-targeted network pharmacology interaction of NK and the formulation. EnrichR was used to depict a sub-network of diabetes proteins and their relationship with diabetes associated comorbidities. NK demonstrated a dose dependent DPP4 inhibition with an IC_50_ of 2.06 μg GAE/mL. Molecular docking identified three compounds namely Terchebin, Locaracemoside B and 1,2,4,6 Tetra o Galloyl Beta D Glucose showing stable interactions with DPP4 similar to the standard drug Vildagliptin. The network pharmacology analysis of NK identified a number of targets like TNFα, TGFβ1, SOD1, SOD2, AKT1, DPP4 and GLP1R in its protein-protein interaction network which are vital to diabetic progression and complications. The present work demonstrated that the polyherbal formulation NK has DPP4 inhibition potential and modulates a large number of diabetes related proteins and pathways. The approach adopted in the current study by combining *in vitro* and *in silico* methods allowed us to understand the mechanism of DPP4 inhibition by the formulation and also the possible pharmacological networking through which the formulation exert its systemic effect in diabetes management.

## 1. Introduction

Dipeptidyl-peptidase IV (DPP4) inhibitors have emerged as an important class of anti-diabetic medication that provide better glycemic control with less adverse effects owing to their role in modulating incretin physiology. One of the major substrates of DPP4 is Glucagon Like Peptide -1 (GLP1), an important incretin hormone secreted by the enteroendocrine cells, known for its multi-organ interactions in regulating body’s glucose homeostasis [1]. The GLP1 mediated incretin effects are highly compromised in type 2 diabetes primarily due to the serine protease action of DPP4 resulting in GLP1 degradation. Consequently DPP4 inhibitors, that can prolong GLP1 action in the body, are considered to be beneficial for improving glucose homeostasis without posing any risk of hypoglycaemia or weight gain [2 - 4]. Five such DPP4 inhibitors namely Sitagliptin (Januvia, Merck), Vildagliptin (Glavus, Novartis), Saxagliptin (Onglyza, Bristol-Myers Squibb), Alogliptin (Nesina, Takeda), and Linagliptin (Tradjenta, Boehringer-Ingelheim) are already in active clinical use and more including Gemigliptin, Teneligliptin, etc. are in the clinical trial stage as well [5-7]. The incretin hormone biology and DPP4 actions are crucial players of the gut centred systemic glucose metabolism of the body (broadly referred as gastrointestinal mediated glucose disposal - GIGD) and therefore targeting DPP4 and incretins become an important strategy for developing more holistic, multi-targeted and systemic drug interventions [5]. Such approaches are imperative for long term management of chronic lifestyle diseases like diabetes and diabetes-associated comorbidities.

Despite its important role in clinical management of systemic glucose homeostasis, the marketed DPP4 inhibitors are reported to have several adverse effects and are relatively expensive [6]. This spurs scientists to look for more affordable and safer substitutes from alternative sources. *Ayurveda* formulations, by virtue of their multicomponent herbal ingredients, form a rich repository of bioactive molecules for screening potential drug candidates [7]. Embracing the *Ayurveda* concept of ‘*Agni*’ (unique concept that describes the biology of metabolism) as well as its gut centric perspective of managing metabolic diseases further enhances the scope of screening *Ayurveda* formulations for identifying potential drug candidates that inhibit DPP4 and thereby modulate incretin biology. In this background, the present study investigates the DPP4 inhibitory effect of an *Ayurveda* polyherbal formulation *Nisakathakadi Kashaya* (NK) prescribed for the clinical management of diabetes and associated comorbidities. The molecular interactions of the key bioactives from NK with DPP4 protein were analyzed using docking and molecular dynamic (MD) simulation tools.

At the frontiers of disease biology, systems and network medicine concepts have begun to redefine the ‘one disease-one target-one drug’ dogma prevalent in current biology [8]. Emerging insights into the mechanism of action of polyherbal formulations also suggests the possibility of herbal formulations (including *Ayurveda* formulations) exerting their overall biological effects through a multitarget and multi-pathway crosstalk [9]. The *Ayurveda* formulation, NK, selected in this study is a combination of eight herbal ingredients with known hypoglycemic effects (Table-1). A variety of phytochemicals identified from these ingredient plants are known for their multitargeted mode of action. Therefore, our study hypothesized that the formulation, besides inhibiting DPP4 enzyme, could also exert a network pharmacology mode of action by interacting with putative genes in the human body [10]. This prompted us to further investigate the spectrum of diabetes related genes that are modulated by the formulation. Network pharmacology and structural bioinformatics tools allowed us to curate and capture the physiologically relevant signalling interactome of multi-component formulation and the pharmacological networking of phytochemicals was deduced via the openly available software tools. A disease-gene overlap analysis performed in the study further provided information on the possible molecular basis for the poly-pharmacological action of NK in diabetes, diabetic complications and diabetes associated comorbidities.

**Table 1:** List of herbs present in *Nisakathakadi Kashaya* (NK)

| S.No | Scientific Name | Common Name | PART USED |
| --- | --- | --- | --- |
| 1 | *Curcuma longa* L. | Haridra, Turmeric | Rhizome |
| 2 | *Strychnos potatorum* L. | Kataka, clearing nut | Seed |
| 3 | *Ixora coccinea* L. | Paranti | Root/Stem |
| 4 | *Symplocos racemosa* Roxb. | Lodhra | Stem bark |
| 5 | *Emblica officinalis* Gaertn. | Amla, Amalaki, Indian Gooseberry | Fruit |
| 6 | *Aerva lanata* (L.) Juss. ex Schult | Gorakshaganja | Entire plant |
| 7 | *Vetiveria zizanioides* (L.) Nash | Ushira, Khas Vetiver | Root |
| 8 | *Salacia reticulata* Wight | Saptachakra, Salacia | Stem/Root |

## 2. Materials and Methods

### 2.1. Reagents and NK quantification

*Nisakathakadi kashaya* (NK) was procured from a leading *Ayurveda* drug manufacturer and the details about the product were recorded in the Sample Data Sheet (SDS) book maintained in the University. The DPP4 enzyme assay kit was procured from Enzo Life Sciences (Cat. No: BML-AK499-0001). In order to quantitatively use the formulation for DPP4 inhibition assay, total tannins present in the NK formulation was estimated using standard Folin - Ciocalteu method using gallic acid as standard. Briefly, 10 μL of the formulation mixed with 40 μL of water, 50 μL Folin’s reagent and 100 μL of 3.5% Na_2_CO_3_ was incubated at room temperature for 30 min. A set of gallic acid standards (50, 25, 12.5, 6.25, 3.125 μg/mL) were prepared in the same manner. The absorbance was measured at 700 nm using a multi-well plate reader (xMark Microplate Spectrophotometer, BioRad, USA). The experimental concentrations of test samples were expressed as ‘μg of gallic acid equivalent tannin (GAE)/mL of sample’

### 2.2. DPP4 inhibition assay

The DPP4 inhibition assay was carried out as per the kit instructions. Briefly, various concentrations of NK were prepared with the assay buffer. The test samples, positive control and the standard inhibitor (given in the kit) were incubated with DPP4 enzyme for 20 minutes at room temperature. The fluorogenic substrate provided with the kit was added and the plate was read for 30 minutes using a fluorometer (Biotek Synergy H1 microplate reader) at Ex:380/EM:460 nm.

The percentage remaining activity in the presence of inhibitor was calculated using the formula,

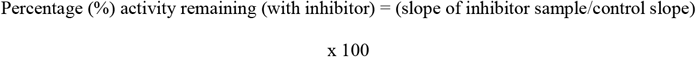

### 2.3. Phytochemical data collection

The phytochemicals present in the eight medicinal plants were extracted from two publicly available databases *viz*. Dr Duke’s Phytochemical and Ethnobotanical Database (https://phytochem.nal.usda.gov) and IMPPAT - Indian Medicinal Plants, Phytochemistry And Therapeutics (https://cb.imsc.res.in/imppat/home) [11]. Phytochemicals reported from the respective parts used in the formulation were shortlisted and considered for further studies. The detailed chemical information about the shortlisted phytochemicals were obtained from the PubChem database (https://pubchem.ncbi.nlm.nih.gov/) and used for molecular docking and other bioinformatics studies.

### 2.4. Preparation of DPP4 structure for docking

The three-dimensional (3D) structure of DPP4 interacted with Vildagliptin (PDB ID: 6B1E) was retrieved from Research Collaboratory for Structural Bioinformatics Protein Data Bank (RCSB PDB; http://www.rcsb.org). The protein was then prepared using the *protein preparation wizard* module available in the Schrodinger suite (Schrödinger, LLC, NewYork, United States, 2020). For acceptable ionisation states, h-bonds corresponding to pH-7 were provided to both basic and acidic amino acid residues followed by energy minimization of the protein structure using the OPLS3e force field. The ligand-binding site of the DPP4 protein was identified from a detailed study of literature. CASTp server was used to validate the binding sites that were found in the literature. The prepared protein generated from the Protein preparation wizard was considered to generate the grid box for the binding of the ligand to the protein molecule. This step was carried out using *the Receptor Grid Generation* module available in Schrodinger maestro Suite. Active site residues of the DPP4 protein (consisting of residues Glu205, Glu206, Asp708, His740, Ser630, Arg125, Ser209, Phe357, Tyr547, Tyr631, Ile651, Trp659, Tyr662, Tyr666, Arg669 and Val711) was selected to define the size of the Grid box. All catalytic active site residues (such as Glu205, Glu206, Asp708, His740 and Ser630) and substrate binding site residues were personally verified and confirmed to be correctly restricted within the rectangular grid box.

### 2.5. Preparation of Ligands

A total of 206 compounds from NK have been shortlisted for the virtual screening of ligands interacting with DPP4. The 3D structures of all compounds in Spatial Data File (SDF) format were downloaded from the PubChem database. *The LigPrep* module of Schrodinger (LigPrep, Schrödinger, LLC, New York, 2021) was used to prepare all the 206 ligands at a pH of 7.0 ± 0.1, and partial atomic charges were applied and ionisation states were developed. All compounds were minimised using OPLS 2005 force field and each ligand was subjected to energy minimization until it passed an RMSD (root mean square deviation) limit of 0.001.

### 2.6. Molecular Docking

The molecular docking of the 206 compounds of NK with DPP4 was done using the XP (extra precision) docking mode of the GLIDE program with default parameters (Glide, Schrödinger, LLC, New York, 2021) [12]. In this process, ligand molecules were treated as flexible and the receptor is considered as rigid entity to attain the most significant interaction with binding site residues of DPP4. Then, using the Prime-MM-GBSA module of the Schrodinger Maestro suite, the binding free energy calculations were performed. Based on the XP-docking score, binding free energy and interaction with the active site residues, top 3 ranked complexes and one protein-known inhibitor (Vildagliptin) complex were considered for further assessment.

### 2.7. Molecular Dynamics (MD) Simulation

The top three bioactive molecules and one known inhibitor (Vildagliptin) complexed with DPP4 protein were subjected to 50ns simulation production run using Desmond (v5.6) package [13]. For each protein-ligand complex, the TIP4P water model was used as solvation medium and the periodic boundary conditions were set to orthorhombic with the size of 10 × 10 × 10 Å. The necessary number of ions was adjusted to neutralise the system and the salt concentration was maintained at 0.01M. To equally distribute the ions and solvent around the protein-ligand complex, each system was equilibrated using the NPT (respectively number of the particle, system pressure and temperature) with a constant temperature of 300K. Using the relaxation model, the simulation production run was carried out for 50ns for all four complexes. Using Simulation Interaction Diagram and Simulation Event Analysis modules of Schrodinger suite, RMSD of the protein backbone, RMSF (Root mean square fluctuation) of individual amino acids and atoms of ligand, RoG of the ligand, SASA of the ligand, intra H-bond calculation of the ligand, etc. were calculated from MD simulation trajectory. The simulation interaction diagram was used to study the interaction details of protein and ligands during the simulation. The thermal_mmgbsa.py script was used to calculate the average binding free energy from the simulation trajectory based on MM-GBSA using a step size of 10.

### 2.8. Target mapping of phytochemicals using network pharmacology tools

The potential target proteins of the phytochemicals were obtained from three database sources *viz*. ChEMBL (https://www.ebi.ac.uk/chembl), STITCH (http://stitch.embl.de/) and BindingDB (https://www.bindingdb.org) [17-19] (Gaulton et al., 2012(Kuhn et al., 2008)Liu et al., 2007). The putative targets with active interaction sources from Experiments and Databases and with the minimum confidence score of 0.400 was used during the STITCH search. The SMILES notation of the compounds was used in BindingDB and the conversion from CID to SMILES was done using the PubChem identifier exchanger (https://pubchem.ncbi.nlm.nih.gov/idexchange/idexchange.cgi). A similarity cut-off of 0.85 was used during the BindingDB search.

### 2.9. Network construction and disease associated gene identification

A phytochemical target protein network was obtained using Cytoscape_v3.9.0 [17]. Every chemical with its representative targets is arranged and files are loaded in Cytoscape. The hub proteins and hub compounds were identified from the network using the Cytoscape plugin cytoHubba [18]. It provides 11 topological analysis methods, out of which MCC is used which captures more essential proteins in the top-ranked list in both high degrees as well as low degrees. MCC indicates that every node is connected to every other node in that subgraph.

### 2.10. EnrichR analysis for identifying target-disease overlap

The reported phytochemical targets from the databases were corrected for duplicate entries and processed for protein disease overlap using the publicly available tool DigiNet (http://www.disgenet.org/web/DisGeNET/), OMIM, and ClinVAR databases containing information about relationships between human/animal genes and proteins and diseases that was accumulated from various sources mainly through text-mining approaches.

The KEGG pathway database and ClueGO plugin in Cytoscape was used for understanding the biological pathways regulated by the potential network constructed for the formulation [19],[20]. A Venn diagram analysis was done to obtain the common targets between diabetic complications and associated metabolic diseases.

## 3. Results

### 3.1. NK inhibits DPP4 dose dependently *in vitro*

NK showed a strong dose dependent inhibition of DPP4 enzyme with an IC_50_ of 2.09μg GAE/mL and a maximum inhibition of 97% at 25μg of GAE/mL and a minimum inhibition of 64% at the lowest concentration of 3.125μg of GAE/mL tested in the study significantly (p<0.001) (Fig-1).

**Figure-1:**
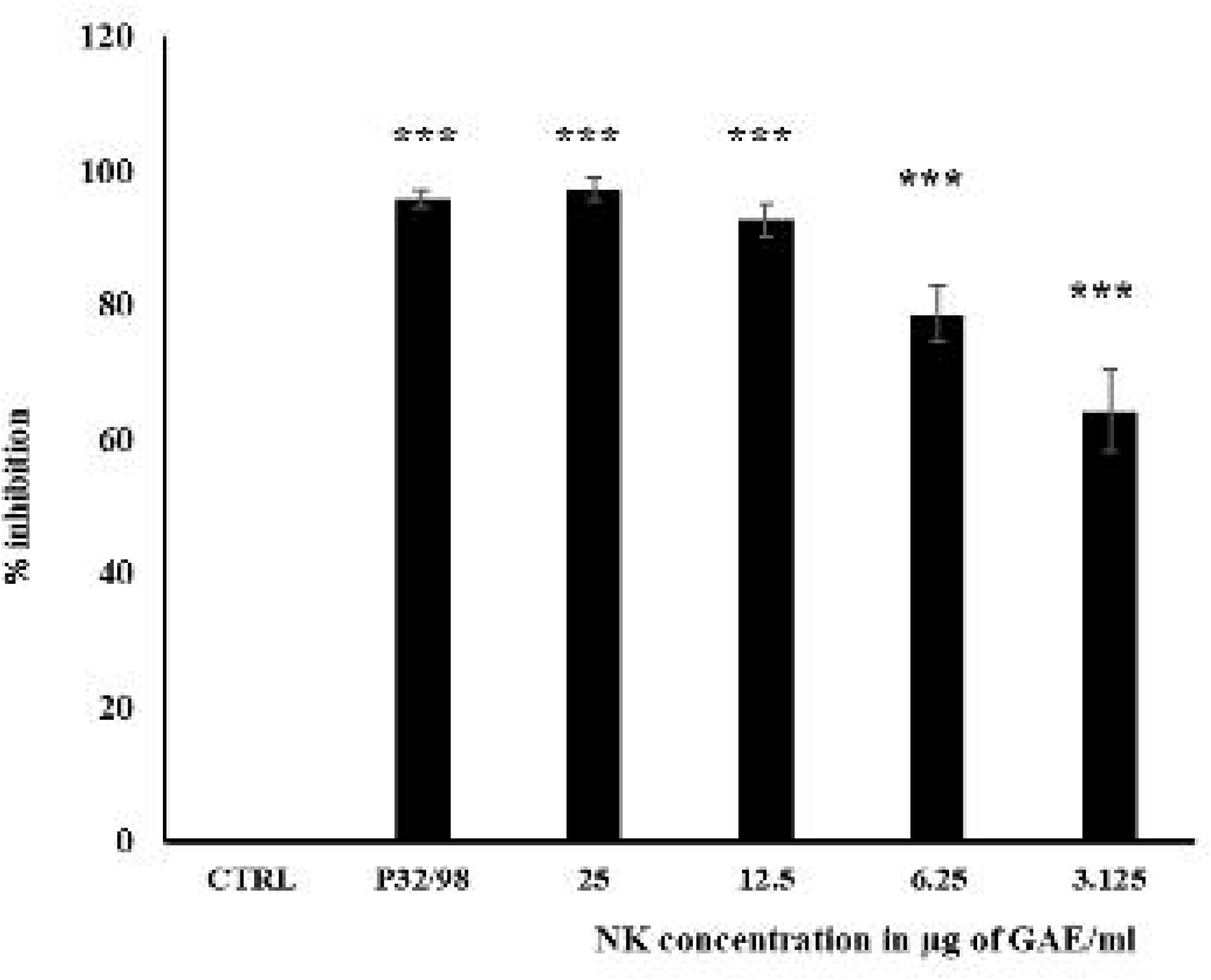
Inhibition of DPP4 by NK formulation. Graph shows a concentration dependent inhibition of DPP4 enzyme action upon treatment with various concentrations of NK (p value ≤ 0.001, ***)

### 3.2. Molecular Docking studies showed key phytochemicals identified in NK directly interacting and inhibiting DPP4 activity

There are 206 phytochemicals identified from eight medicinal plants present in NK. The list of herbs present in NK are listed in Table-1. In order to identify the phytoconstituents of NK responsible for DPP4 inhibition, a virtual screening of all the 206 compounds were performed by docking them to DPP4 protein. Well known DPP4 inhibitor Vildagliptin was used as a positive control. The top-ranked 35 compounds, based on their docking score, and MM-GBSA are listed in Table-2. The docking and MM/GBSA scores for Vildagliptin were -3.899 kcal/mol and -26.7 kcal/mol, respectively. The compounds listed in Table 2 showed a better docking score compared to that of Vildagliptin. The interaction details of the top 3 compounds from the table *viz*. Terchebin, Locoracemoside B and 1,2,4,6 Tetra o Galloyl Beta D Glucose (TGBG) having G binding energy -47.12 kcal/mol, -45.45kcal/mol, and -60.73 kcal/mol, respectively, are reported in the study along with Vildagliptin as a control. DPP4 amino acid residues Glu206, Tyr666 and Asn710 that are involved in Vildagliptin-DPP4 interaction are found to interact with the top 3 compounds identified from the virtual screening (Fig-2). Out of the 35 compounds, MM/GBSA scores were not obtained for Kotalanol and Corilagin.

**Table 2:**
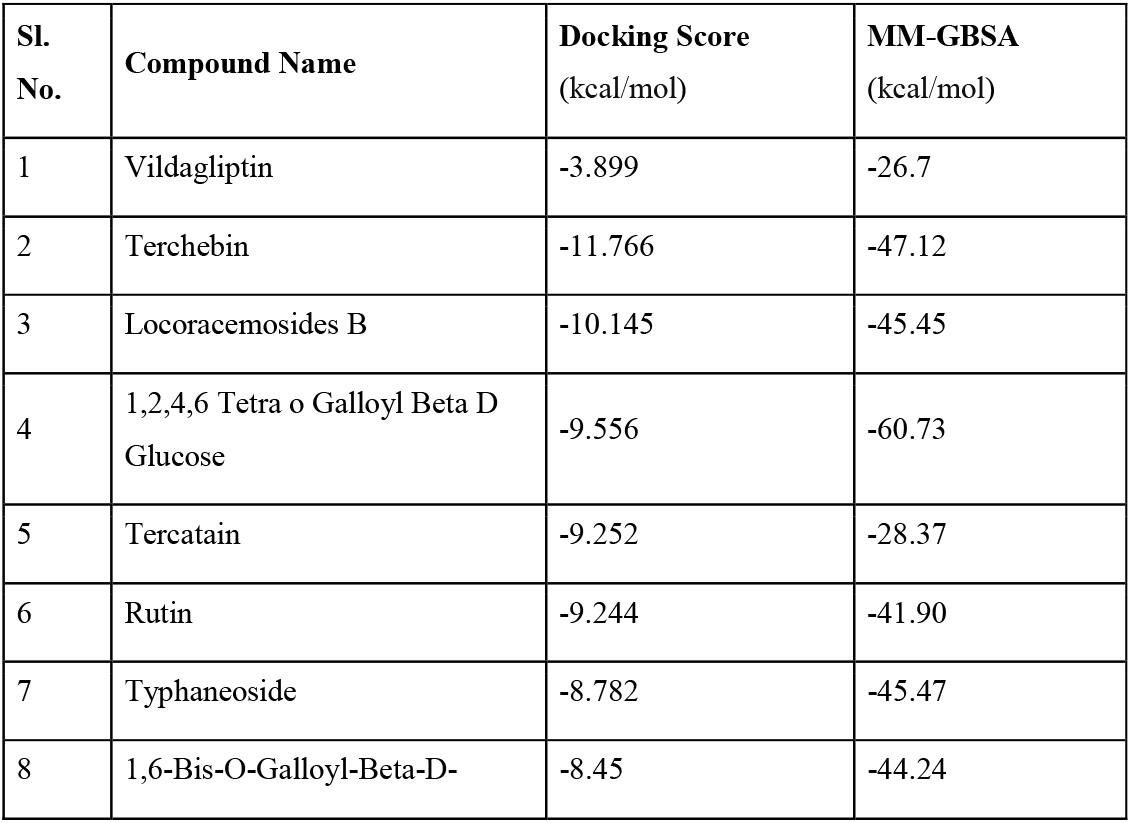
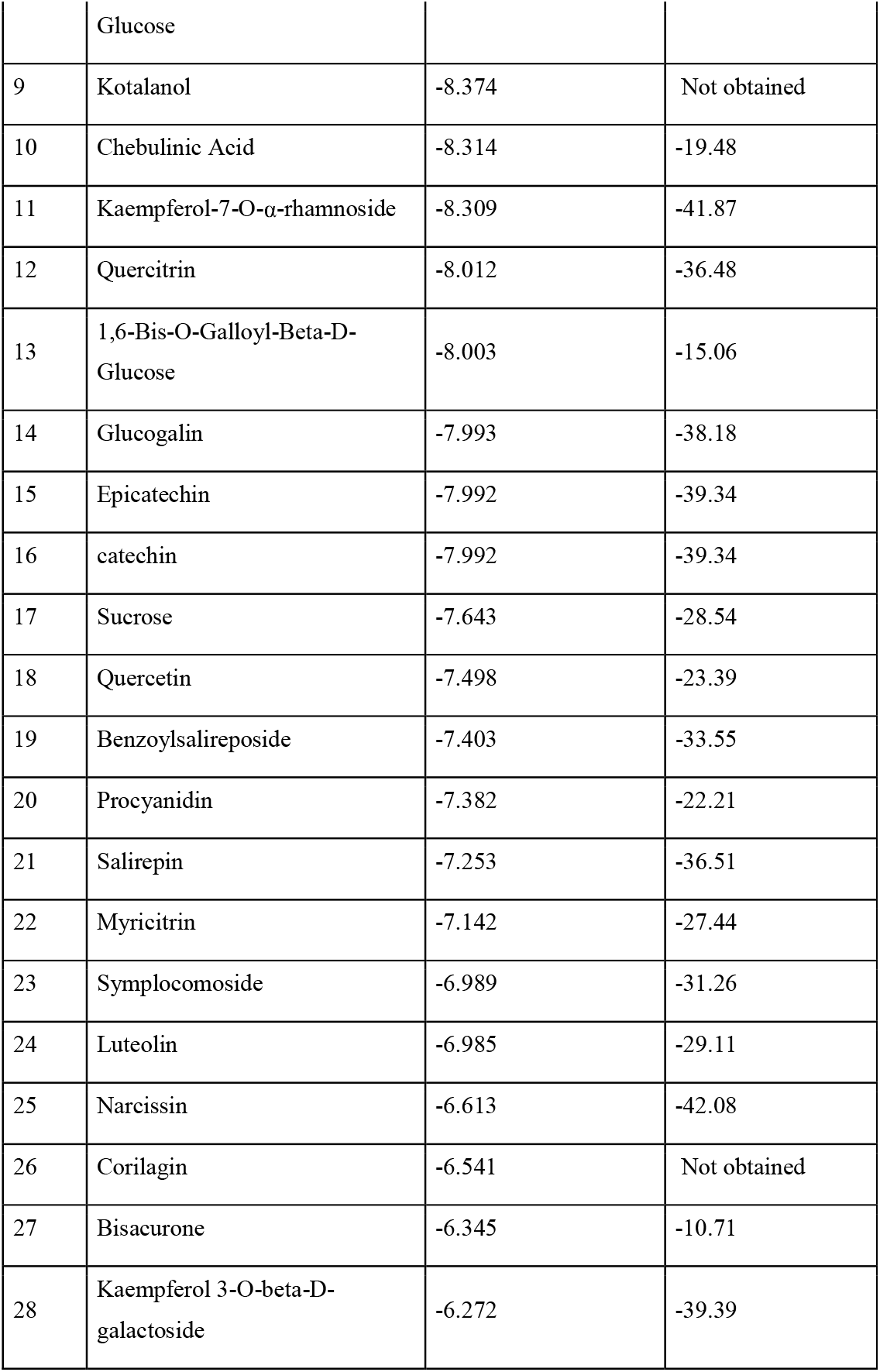
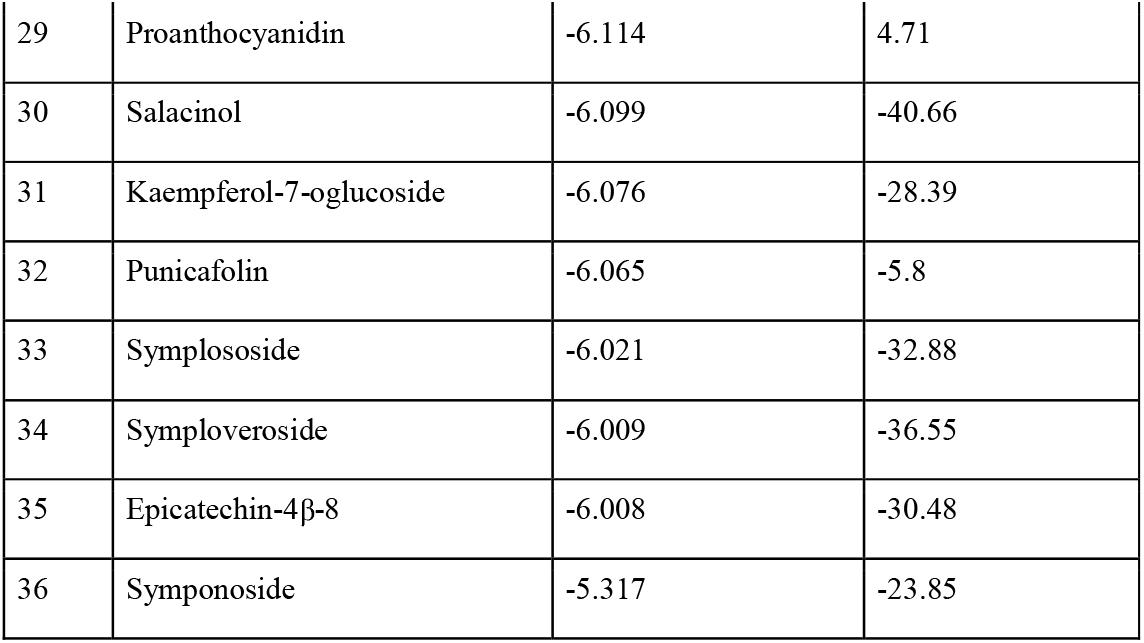
Top scored 35 compounds from the Virtual Screening of NK phytochemicals with DPP4. The docking score and binding free energy from MM-GBSA analysis are reported

**Figure-2:**
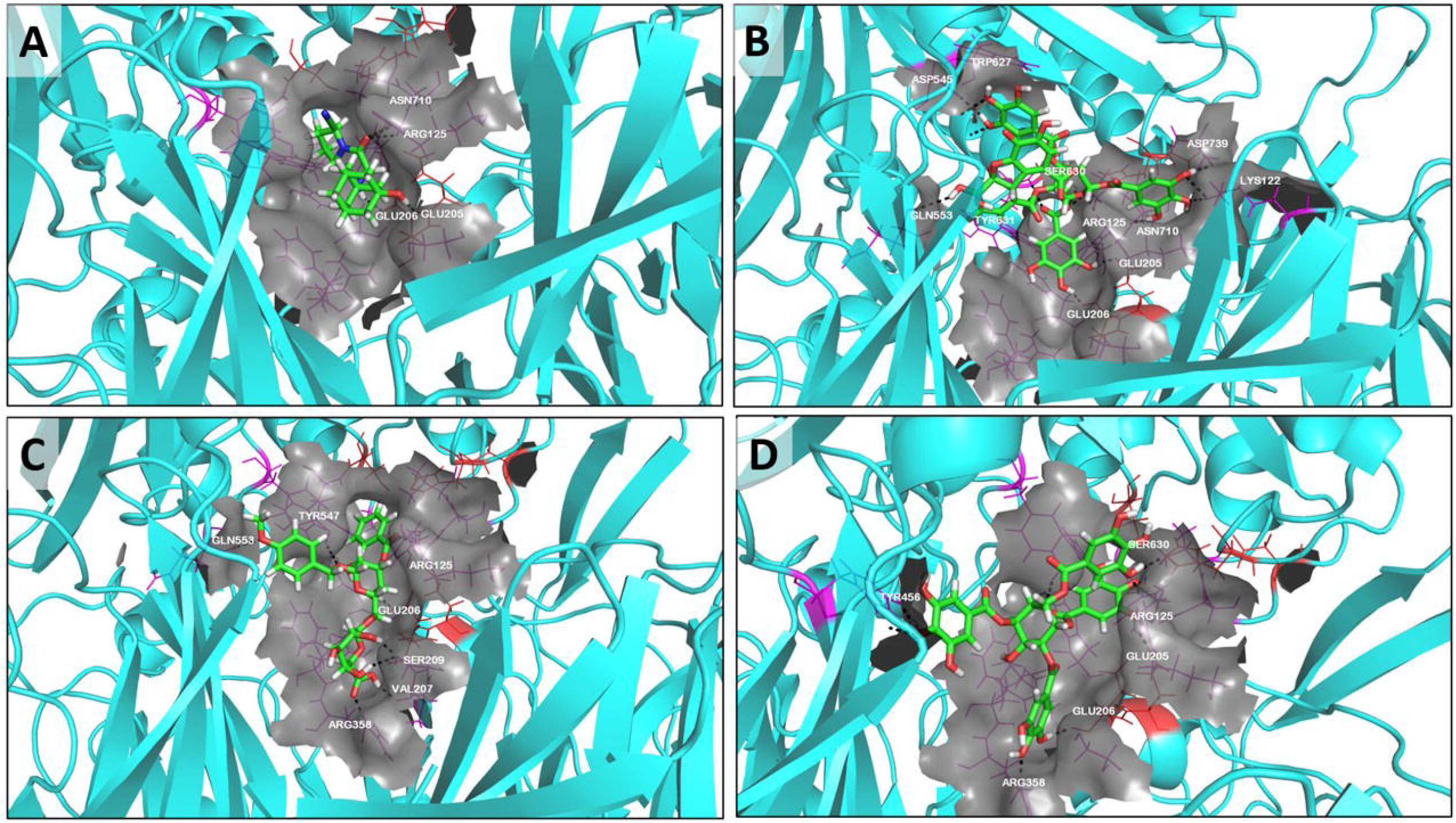
Interaction of DPP4 with various ligands. Interactions of Vildagliptin (A), Terchebin (B), Locoracemoside B (C) and 1,2,4,6 Tetra o Galloyl Beta D Glucose (D) with DPP4.

### 3.3. MD simulation analysis of DPP4 ligand complex

The top three compounds selected from the docking study, Terchebin, Locoracemoside B, and TGBG as well as Vildagliptin were further subjected to a detailed interaction analysis using MD simulation. MD is a powerful computer simulation technique that is now being investigated in the field of computer-aided drug discovery research to better understand how the protein-ligand complex behaves in a dynamic environment at the atomic level over a user-specified time period. A 50ns MD simulation analysis of the DPP4-ligand docked complex was performed and the RMSD profiles of the protein and ligands during the 50ns simulations are analysed (Fig-3). The average RMSD of DPP4 and Vildagliptin was found to be 1.75Å and 2.0Å, respectively (Fig-3A). Whereas, the average RMSD of DPP4 and Terchebin was 2.8Å and 3.5Å; DPP4 and Locoracemoside B was 2.1Å and 1.5Å and DPP4 and 1,2,4,6 Tetra o Galloyl Beta D Glucose was 2.1Å and 1.8Å, respectively (Fig-3B-D). All ligands were found to have stable interactions in the active site.

**Figure-3:**
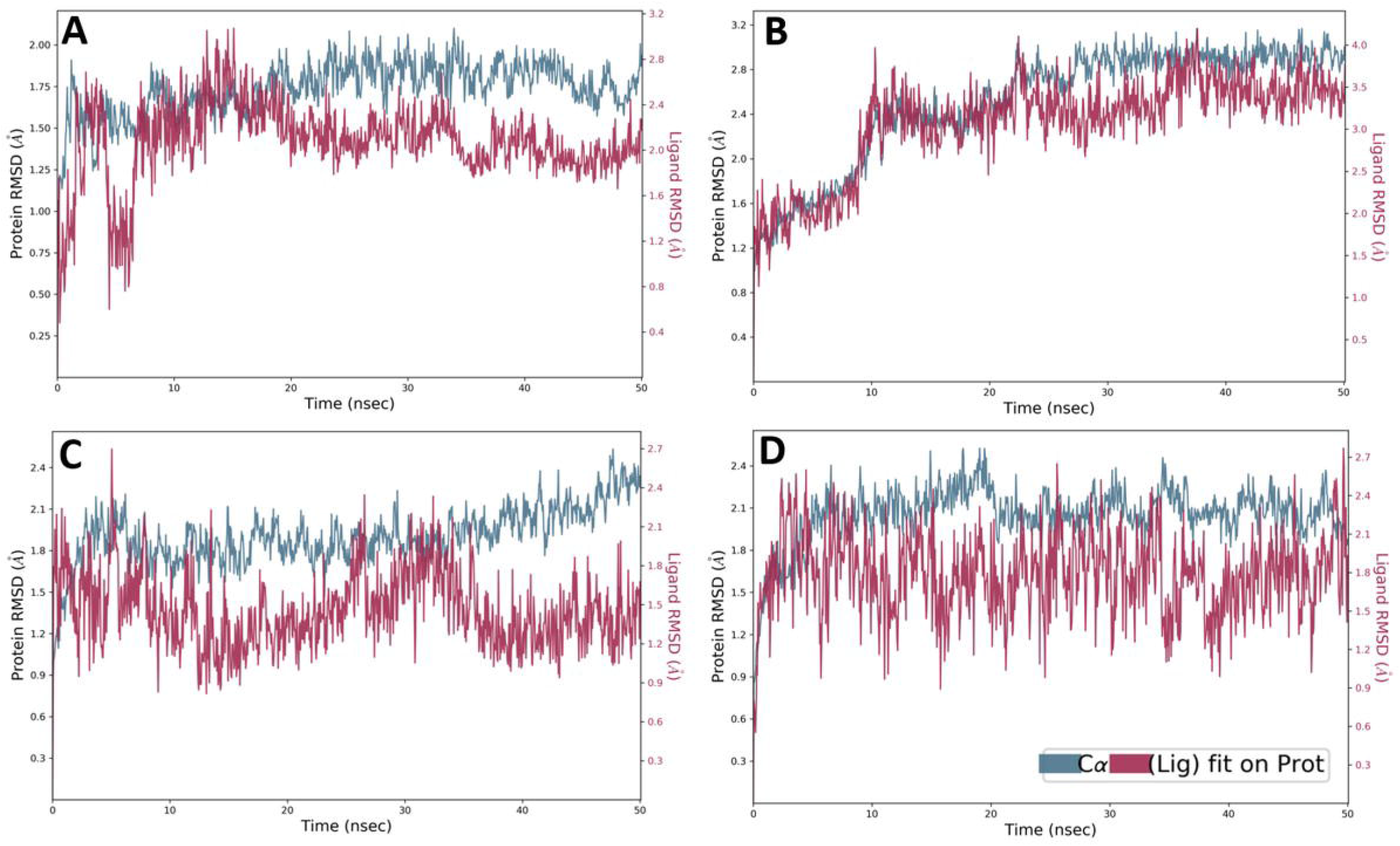
RMSD plot. RMSD plot of Protein and Vildagliptin (A); Protein and Terchebin (B); Protein and Locoracemoside B (C); Protein and 1,2,4,6 Tetra o Galloyl Beta D Glucose (D).

### 3.4. Stability analysis of phytochemicals interacting with DPP4

The protein-ligand interactions are depicted for 4 interactions (hydrogen bond, hydrophobic bond, water bridge and ionic bond) using the clubbed bar graph. The interactions with the specific amino acids throughout the run are depicted using normalised values. The stability is majorly defined by the hydrogen bonds being formed between the ligand and the protein. The DPP4-Vildagliptin interactions showed 4 hydrogen bonds being formed throughout the run at positions Glu205, Glu206 and His740. Strong hydrophobic interactions were seen at positions Val656 and Tyr666 (Fig-4A). For DPP4-Terchebin interaction, 8 hydrogen bonds at positions Ser206, Ser242, Tyr545, Lys554, Arg560, Trp627, Asn710 and Asp739 as well as hydrophobic interactions at positions Lys122, Arg125, Tyr547, Trp627 and Trp629 were identified (Fig-4B). Similarly, a total of 10 hydrogen bonds at positions Glu206, Val207, Ser209, Arg358, Tyr547, Ser552, Gln553, Tyr385 and Ser630 and hydrophobic bonds at Phe357, Tyr547, Tyr585, Tyr631, Tyr662, Tyr666 and Val711 were found in the DPP4-Locoracemosids B interaction (Fig-4C). The DPP4-TGBG interaction showed hydrogen bonds at Glu205, Glu206, Val207, Ser209, Tyr547, Pro550, Tyr662, Arg669, Tyr670 and Asn710 positions and hydrophobic bonds at positions Phe357, Tyr547, and Tyr666 (Fig-4D). The trajectory was subjected to detailed binding energy analysis using thermal_mmgsa script. Free energy of binding for Vildagliptin, Terchebin, Locoracemoside B and TGBG found to be -28.6 kcal/mol, -66.5kcal/mol, -58kcal/mol and -90kcal/mol, respectively.

**Figure-4:**
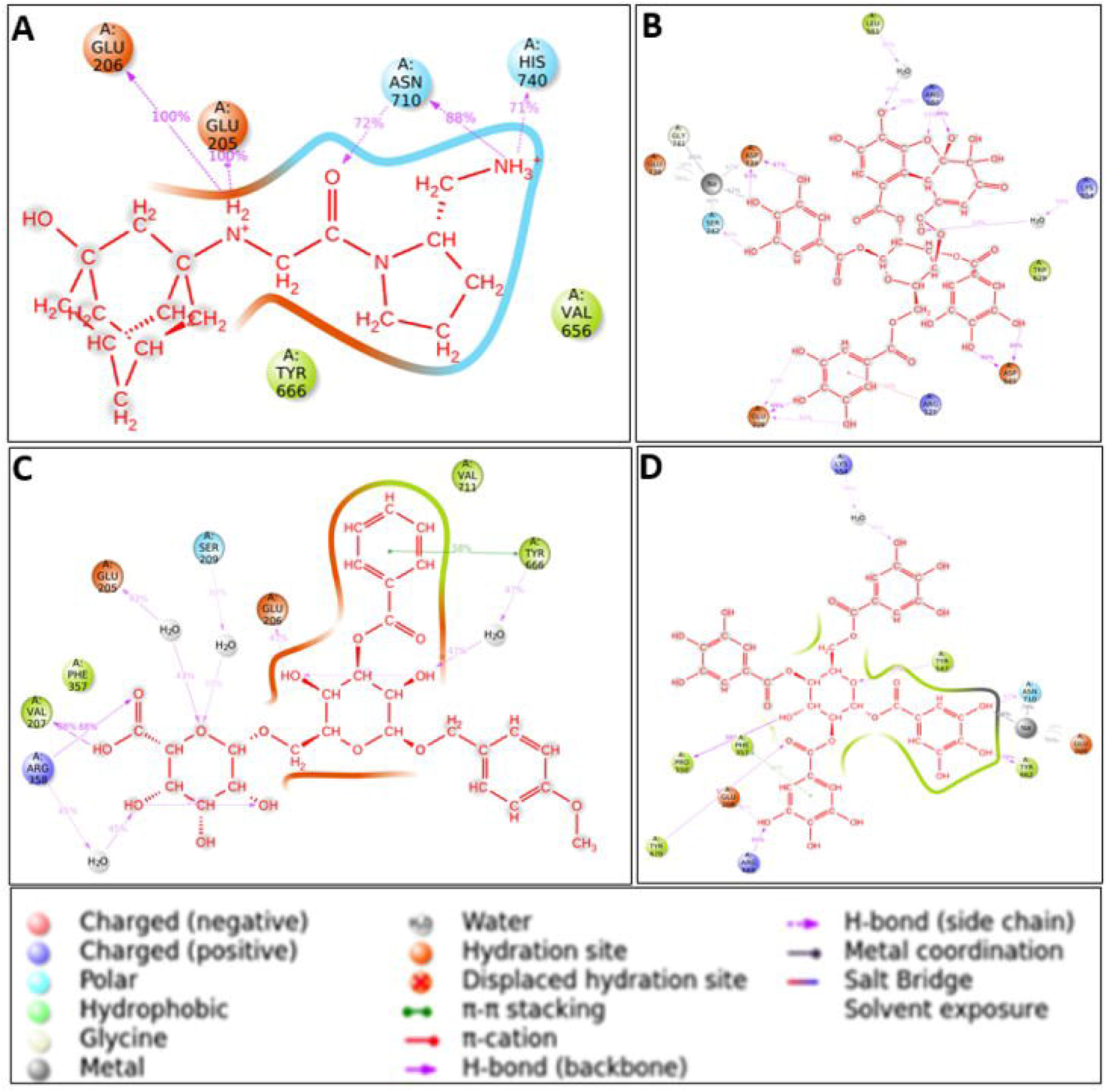
2D interaction diagram of ligands with protein during the simulation. (A)-Vildagliptin, (B)-Terchebin, (C)-Locoracemoside B and (D)-1,2,4,6 Tetra o Galloyl Beta D Glucose with DPP4.

### 3.5. Target mapping of phytochemicals identified in NK formulation for network analysis

The constituent bioactives in a herbal formulation can interact with diverse bioactives to bring out their systemic biological effects and this can be studied using network pharmacology analysis methods, enabling us to delineate the possible mode of action of the formulation and hypothesise new biological pathways and interactomes involved in its pharmacological interactions [10].

The targets of 206 phytochemicals were curated from three databases as mentioned in materials and methods viz, STITCH, ChEMBL and BindingDB. Out of 206 bioactives studied, only 139 bioactives were found to have reported protein targets in these three databases and a total of 1555 proteins were identified for these 139 compounds. The remaining 67 compounds were found to have no interaction reports in the top databases widely used for target identification.

Using the Cystoscape software, the phytochemicals and their identified targets were then converted to a compound-target network to represent the possible biological crosstalk modulated by NK. The network has 1694 nodes and 3264 edges identified (Fig-5A). The compound-target network is a bipartite one, representing the interaction between phytochemicals and their putative targets. From this network, we observed that curcumin has the maximum interactions with 138 targets. Several hub compounds like chrysin and rutin which showed large interactions with proteins in the network were also reported to inhibit DPP4 *in vitro* [20],[21]. The list of all the phytochemicals and their targets can be found in Supplementary Data 1. The complete list of hub proteins and hub compounds are given in Table 3 and 4 (abbreviations can be accessed from Supplementary Data 6). (The network pharmacology studies also revealed compounds such as rutin, typhaneoside, myricitrin, narcissin, kaempferol 3-O-beta-D-galactoside and quercitrin having DPP4 as their targets is aligning with our docking results which showed strong DPP4 inhibition by these compounds.

**Figure-5:**
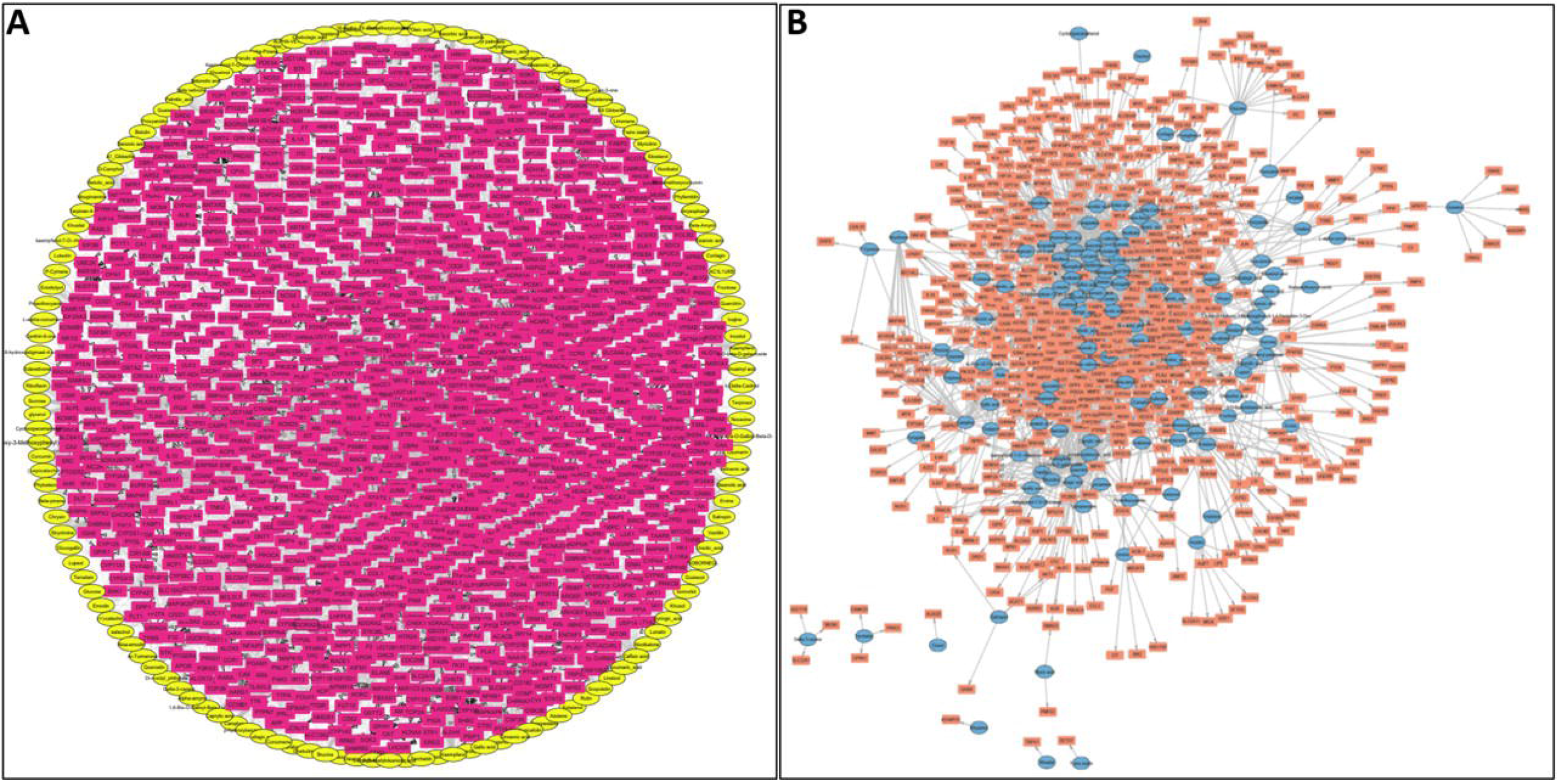
The compound-target network of NK. (A) – The yellow nodes represent the compounds from plants used in the NK formulation and the pink nodes represent the target proteins. (B) -The network of phytochemicals of NK and diabetes-related proteins. The blue color nodes represent the compounds and the orange nodes represent the diabetes associated target proteins.

**Table 3:** List of hub proteins ranked as per their degree values

| S. No . | Hub Gene | Degree | S. No . | Hub Gene | Degree | S. No. | Hub Gene | Degree | S. No. | Hub Gene | Degree |
| --- | --- | --- | --- | --- | --- | --- | --- | --- | --- | --- | --- |
| 1 | CA2 | 26 | 38 | DHCR24 | 8 | 75 | CNR2 | 6 | 112 | GANC | 5 |
| 2 | CYP19A1 | 22 | 39 | AKR1B10 | 8 | 76 | CHRM4 | 6 | 113 | ALOX12 | 5 |
| 3 | CA7 | 18 | 40 | TOP2A | 8 | 77 | AGTR2 | 6 | 114 | PLAT | 5 |
| 4 | ESR1 | 17 | 41 | ALB | 8 | 78 | PNLIP | 6 | 115 | TP53 | 5 |
| 5 | F2 | 16 | 42 | ADRA2B | 8 | 79 | CASP8 | 6 | 116 | PYGL | 5 |
| 6 | AKR1B1 | 16 | 43 | CYP1A2 | 8 | 80 | BCL2 | 6 | 117 | SULT1E1 | 5 |
| 7 | CA12 | 16 | 44 | ALOX5 | 8 | 81 | UGT2B15 | 6 | 118 | CSNK2A1 | 5 |
| 8 | AR | 15 | 45 | PSMB5 | 8 | 82 | TYR | 6 | 119 | GSK3B | 5 |
| 9 | CASP3 | 15 | 46 | NR1I2 | 8 | 83 | XDH | 6 | 120 | SI | 5 |
| 10 | ACHE | 15 | 47 | CA1 | 8 | 84 | GLO1 | 6 | 121 | TTR | 5 |
| 11 | PTPN1 | 13 | 48 | GAA | 8 | 85 | NR3C2 | 6 | 122 | AKR1A1 | 5 |
| 12 | RORC | 13 | 49 | CHRM5 | 8 | 86 | SLC2A1 | 6 | 123 | FAAH | 5 |
| 13 | SHBG | 13 | 50 | HMGCR | 7 | 87 | HDAC6 | 6 | 124 | GSTM3 | 5 |
| 14 | MMP9 | 13 | 51 | GRIN2B | 7 | 88 | SERPINE<br>1 | 6 | 125 | LOX | 5 |
| 15 | ESR2 | 12 | 52 | ITGB3 | 7 | 89 | PTPN2 | 6 | 126 | DHODH | 5 |
| 16 | NR1H3 | 12 | 53 | MAPK3 | 7 | 90 | PLAU | 6 | 127 | CYP3A4 | 5 |
| 17 | UGT1A7 | 12 | 54 | CASP7 | 7 | 91 | CYP51A1 | 6 | 128 | VEGFA | 5 |
| 18 | UGT1A8 | 12 | 55 | F7 | 7 | 92 | MGLL | 6 | 129 | PSMB1 | 5 |
| 19 | UGT1A1<br>0 | 12 | 56 | RGS4 | 7 | 93 | AMY2A | 6 | 130 | CACNA<br>1H | 5 |
| 20 | CA4 | 12 | 57 | ADRA2<br>C | 7 | 94 | RPS6KA3 | 6 | 133 | COMT | 5 |
| 21 | CYP17A<br>1 | 11 | 58 | CASR | 7 | 95 | CA14 | 6 | 134 | MGAM | 5 |
| 22 | CYP1A1 | 11 | 59 | GLA | 7 | 96 | BACE1 | 6 | 135 | PLAUR | 5 |
| 23 | CRYAB | 10 | 60 | PDE5A | 7 | 97 | CTSD | 5 | 136 | IMPA1 | 5 |
| 24 | EGFR | 10 | 61 | CASP9 | 7 | 98 | CYP142 | 5 | 137 | EBP | 5 |
| 25 | TOP1 | 10 | 62 | UGT1A1 | 7 | 99 | ITGAV | 5 | 138 | LSS | 5 |
| 26 | F10 | 10 | 63 | CYP1B1 | 7 | 100 | GRIN1 | 5 | 139 | TNF | 5 |
| 27 | HSD11B<br>1 | 10 | 64 | NOX4 | 7 | 101 | CYP7A1 | 5 | 140 | AVPR1A | 5 |
| 28 | PTGS2 | 9 | 65 | LCK | 7 | 102 | MPO | 5 | 141 | SLCO1B<br>3 | 5 |
| 29 | NR1H2 | 9 | 66 | KDM1A | 7 | 103 | CRABP2 | 5 | 142 | APP | 5 |
| 30 | MAPK1 | 9 | 67 | SQLE | 7 | 104 | GNB1 | 5 | 143 | CYSLTR<br>1 | 5 |
| 31 | AKT1 | 9 | 68 | GRIN2A | 6 | 105 | HTR1B | 5 | 144 | EDNRB | 5 |
| 32 | ADRA2<br>A | 9 | 69 | OSBP2 | 6 | 106 | LPAR3 | 5 | 145 | FABP4 | 5 |
| 33 | UGT1A9 | 9 | 70 | SREBF2 | 6 | 107 | MTRNR2<br>L2 | 5 | 146 | FABP5 | 5 |
| 34 | MAOA | 9 | 71 | CYP2B6 | 6 | 108 | HCAR3 | 5 | 147 | FFAR1 | 5 |
| 35 | DPP4 | 9 | 72 | MAPK8 | 6 | 109 | TAS2R40 | 5 | 148 | NOS2 | 5 |
| 36 | UGT1A3 | 9 | 73 | BCHE | 6 | 110 | TAS2R43 | 5 |  |  |  |
| 37 | MMP2 | 9 | 74 | LPAR2 | 6 | 111 | P2RY4 | 5 |  |  |  |

**Table 4:** List of hub compounds ranked as per their degree values

| S.No | Hub Compound | Degree |
| --- | --- | --- |
| 1 | Curcumin | 138 |
| 2 | Chrysin | 121 |
| 3 | Kaempferol | 106 |
| 4 | Oleic Acid | 87 |
| 5 | Quercetin | 82 |
| 6 | Stearic_Acid | 80 |
| 7 | Caffeic Acid | 78 |
| 8 | Camphor | 78 |
| 9 | Ursolic_Acid | 75 |
| 10 | Ascorbic Acid | 75 |
| 11 | Sucrose | 74 |
| 12 | Oleanolic Acid | 67 |
| 13 | Palmitic_Acid | 67 |
| 14 | Ellagic Acid | 61 |
| 15 | Quercitrin | 60 |
| 16 | Kaempferol 3-O-Beta-D-Galactoside | 58 |
| 17 | Strychnine | 56 |
| 18 | Cinnamic Acid | 55 |

In order to understand the significant pathways associated with these proteins, a KEGG pathway analysis was done. The analysis revealed a number of significant (p-value < 0.05) diabetes associated pathways like PI3K-Akt signalling pathway, MAPK signalling pathway, lipid and atherosclerosis, Ras signalling pathway, chemokine signalling pathway, diabetic cardiomyopathy, insulin signalling pathway, cellular senescence, HIF-1 signalling pathway, AGE-RAGE signalling pathway in diabetic complications and insulin resistance are associated with these proteins.

Also, a ClueGO analysis performed to identify the potential biological processes that are regulated by the formulation showed several metabolic processes are associated with these identified target proteins (Fig-6). The complete list of pathways and processes is given in Supplementary Data 2. These observations support the *Ayurveda* rationale of using NK for the management of diabetic associated symptoms and complications.

**Figure-6:**
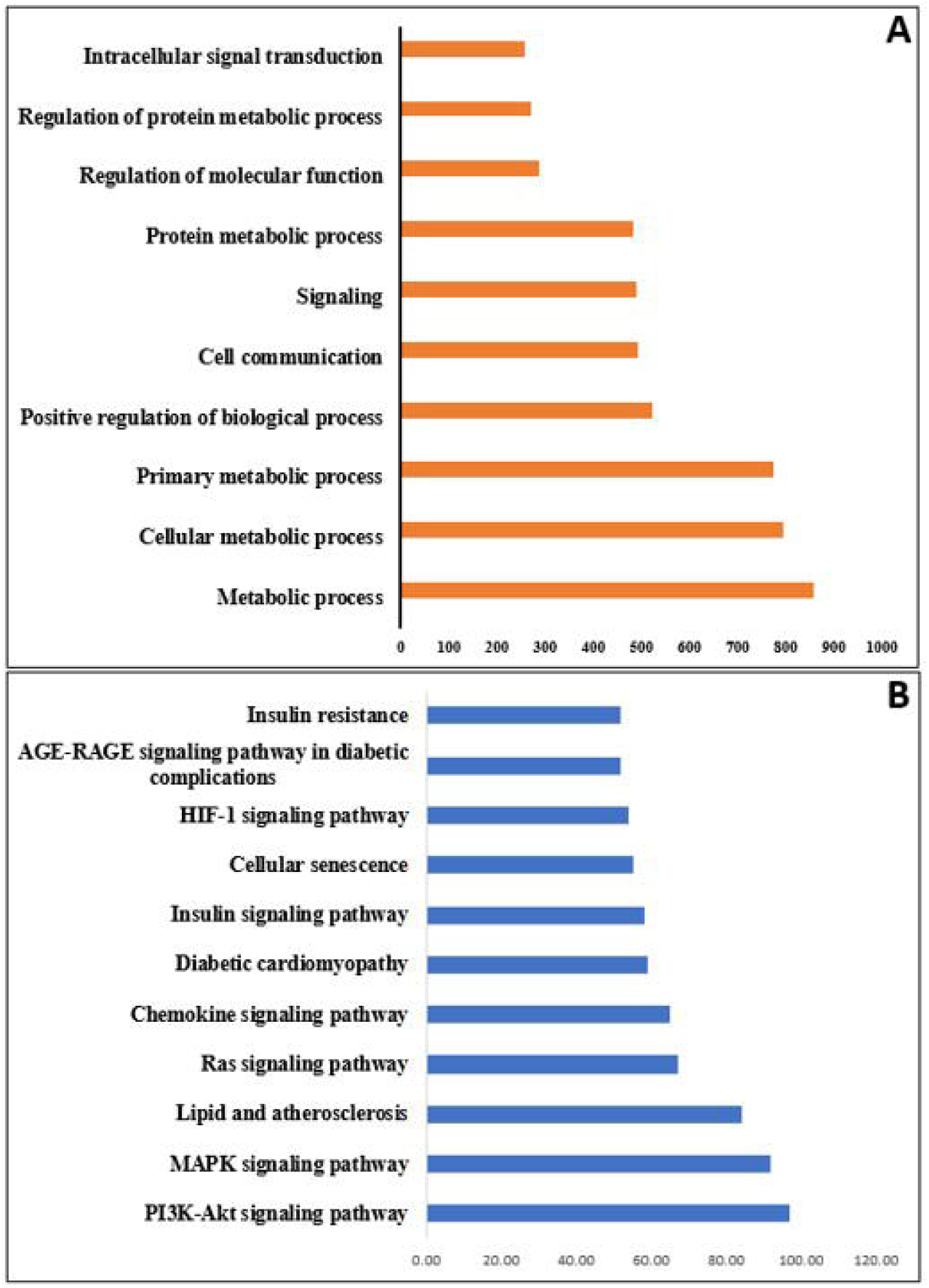
ClueGO analysis of the 1555 target proteins. (A) analysis of the 1555 target proteins involved in different biological processes and (B) analysis of the 1555 target proteins involved in different pathway analysis

### 3.6. Disease overlap analysis of the protein targets mapped for phytochemicals in NK formulation

To get a better idea on the disease associated implications of NK, we used the openly accessible tool EnrichR, a comprehensive gene enrichment analysis tool containing diverse gene set libraries available for analysis [22]. The EnrichR tool was used for disease analysis of the 1555 targets mapped (Supplementary data 5), and data was extracted from three sources viz, DigiNet, ClinVar and OMIM expanded (Supplementary data 5). A total of 9000 disease indications emerged from EnrichR analysis, for which a p-value of ≤ 0.05 was applied to short list the diseases. Further, from the entire list of disease conditions, we used the key words “Diabetes mellitus”, “Insulin resistance”, “Hyperglycemia”, “Hypoglycemia”, to shortlist 614 diabetes associated proteins. (Supplementary Data 3). From this, a subnetwork of diabetes specific proteins was created which contained 743 nodes and 1612 edges (Fig-5B). This showed the putative diabetic interactome of NK comprising a number of critical regulators in diabetes related complications and comorbidities.

For investigating the association of NK targets in diabetic complications and diabetes associated risk conditions, the 614 proteins were again subjected to EnrichR analysis. The resulting disease analysis showed various diabetic complications like nephropathy, neuropathy, cardiomyopathy and retinopathy. and also revealed the presence of related conditions such as fatty liver, obesity and general metabolic disturbances along with inflammation which is a characteristic of all these diseases [23]. We have grouped these conditions into 5 specific categories viz; fatty liver, obesity, general metabolic disturbances, inflammation and diabetic complications. The details of these categories and the complete list of indications are given in supplementary data (Supplementary Data 3 and 4).

To gain an insight into the gene association of diabetes with these 5 categories, we performed a Venn diagram analysis. The results of Venn analysis as observed (Fig-7A) showed that 37 proteins were common among all the diabetes associated diseases and diabetic complications. Proinflammatory markers like NFKB1 and TNFα, characteristic of the insulin resistance were present in the cluster along with lipid regulatory and obesity-linked PPARα and PPARγ [26-28] The other critical markers were insulin signalling molecules like AKT1 and PI3K, demonstrating that these core pathways involved in diabetic pathogenesis can be regulated by multitargeted action of NK [26]. One of the overlap categories in the Venn analysis is obesity, fatty liver and diabetic complications which has 27 shared proteins. We observed that both DPP4 and GLP1R were present in the cluster which aligned with the recent evidence from various studies establishing DPP4 as a crucial gene in the regulation of fatty liver, obesity and diabetic complications. [27].

**Figure-7:**
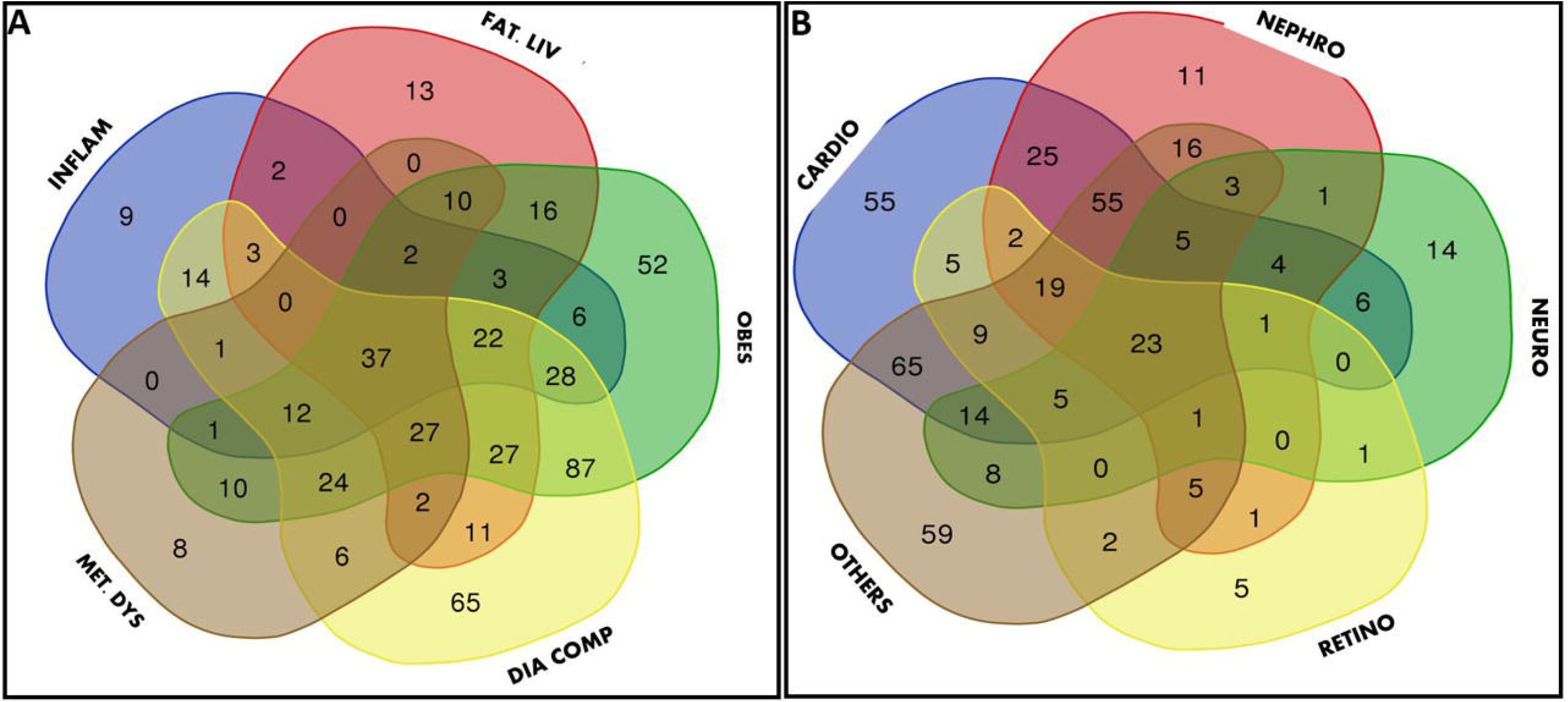
Venn diagram analysis. (A) represents the analysis of protein association between diabetic complications and associated diseases (B) shows the Venn analysis of proteins within diabetic complications. Abbreviations: INFLAM – Inflammation; FAT LIV - Fatty liver; MET DYS - Metabolic disturbances; DIA COMP - Diabetic complications; CARDIO – Cardiomyopathy; NEPHRO – Nephropathy; NEURO – Neuropathy; RETINO – Retinopathy; OTHER - Other diabetic associated complications like diabetic foot disorder, delayed wound healing, microvascular and macrovascular complications

For a specific analysis of the possible overlap within diabetic complications category we further grouped the ‘diabetic complications’ into 5 groups viz. diabetic retinopathy, cardiomyopathy, nephropathy and neuropathy and ‘other diabetic complications. From this specific diabetic complications Venn analysis, a cluster of 23 proteins were identified as common (Fig-7B) and included several important markers such as TNFα, TLR4, CCL2, SOD1, TGFβ1 and SOD2, all of which are deregulated and involved in various stages of diabetic pathogenesis [25],[30,31]. The proteins present in each category and Venn diagram overlaps are listed in Supplementary Data 4.

From our *in silico* studies, we observed that the bioactives in NK can target multiple proteins and pathways which are therapeutically relevant for diabetes and associated complications. We hypothesize that NK with its network interaction can potentially help in managing metabolic disorders. Briefly our *in vitro* and *in silico* analyses, indicates that one of the anti-diabetic mechanisms of actions of NK could be primarily through DPP4 inhibition, thereby modulating GLP1 levels.

## 4. Discussion

DPP4 has gained importance as a key therapeutic target over the last few years due to its primary role in increasing endogenous GLP-1 levels, which is found to be important for improving glycemic control and systemic glucose homeostasis. Though there are several approved DPP4 inhibitors, they are relatively expensive and are associated with some side effects such as pancreatitis, joint pain and increased risk of cancer[6], [32-34]. Hence there is an increasing interest in identifying new compounds that are cheap, safe and accessible.

*Ayurveda* formulations and their component plants, by virtue of their long-established clinical efficacy and multi-component-multi-target mode of action, have been a source for identifying novel bioactives with better efficacy and potentially fewer side effects [7]. Furthermore, the concepts of *Ayurveda* have been a treasure for science-curious people to unearth new biological underpinnings of health and disease manifestations. The *Ayurveda* strategy of holistic healing and the use of poly-herbal preparations have gained global interest and are being scientifically explored through evidence-based research for better understanding of their biological actions [32]. The present study is an attempt to evaluate the poly-pharmacological actions of a clinically established formulation NK with a focus on its ability to inhibit DPP4, one of the relevant targets for diabetes and a potential target for many other complex metabolic diseases. As per the Ayurveda pharmacology, the herbal ingredients of this formulation have the potential to manage *prameha* (≈ diabetes) as well as alleviate disease conditions like *vrana* (wounds) and *netra roga* (eye disorder like retinopathy and cataract) which are clinically considered as diabetic complications [33]. Therefore, Ayurveda physicians prescribe NK for patients showing manifestations of diabetes and diabetes associated complications. Various *in vitro, in vivo* and clinical research evidences support the hypoglycemic potential of the component plants of NK *viz. Curcuma longa, Emblica officinalis, Salacia reticulata, Symplocos racemosa, Vetiveria zizanioides, Strychnos potatorum* and *Aerva lanata* [36*-*41]. The Ayurveda indications of NK and biomedical research evidences of its constituent plants make it an attractive candidate for Ayurveda biology research.

In our study, *in vitro* DPP4 inhibition analysis demonstrated a strong inhibition of the enzyme with an IC_50_ of 2.06 μg GAE/mL, suggesting a beneficial role of NK in improving the incretin biology. Parallel studies conducted in our laboratory have also shown that NK increases GLP1 secretion from enteroendocrine cells (unpublished data). It is well-known that Ayurveda formulations will have multiple bioactive compounds that can act on targets individually or in a synergistic manner. Although the formulation as a whole is shown to inhibit DPP4 it is important to know the nature and mode of action of bioactive(s) responsible for the inhibition. Computational methods are used for predicting and studying the mechanisms of traditional medicinal formulations to circumvent the challenges associated with classical drug discovery.[36]. With this focus, an *in silico* data mining and virtual screening of phytochemicals present in NK was carried out for molecular docking and computational biology analyses. The results showed three compounds Terchebin, Locoracemoside B and TGBG having strong affinity for DPP4 and stable interactions comparable to the standard inhibitor Vildagliptin used in our study. The DPP4 protein consists of Glu205, Glu206, Asp708, His740 and Ser630 residues that are essential for Vildagliptin binding and inhibition. In addition to these, other residues such as Arg125, Ser209, Phe357, Tyr547, Tyr631, Ile651, Trp659, Tyr662, Tyr666, Arg669, and Val711 are also found to play important role in protein-ligand binding. Both Vildagliptin (standard inhibitor) and the phytochemicals datamined from NK virtual screening are found to interact with same residues such as Glu205, Glu206, Tyr666 and Asn710 that are involved in DPP4 inhibitor interaction. This validates our in vitro DPP4 inhibition experiment and provides a possible mechanism of action through which the formulation inhibits DPP4. Terchebin and TGBG are reported from *Emblica officinalis*, which is considered as a ‘wonder plant’ in *Ayurveda* and it is widely reported for its antidiabetic and other biological actions [37],[38]. Locoracemoside B is a glycoside isolated from *Symplocos racemosa*, which is another important plant grouped under the anti-obesity (*medo-hara*) groups of plants in Ayurveda,[39].

Apart from being a source of structurally diverse group of molecules, polyherbal preparations are expected to have a multi-targeted pharmacological networking of phytocompounds to exert their biological action. In one of the landmark papers on network pharmacology, Hopkins *et al* emphasized the relevance of this poly-pharmacology in overcoming the challenges of complex disease management [40]. He suggested that complex diseases may require a multi targeted therapeutic strategy for getting the desired results. This concept of pharmacological networking is much relevant in the context of Ayurveda formulations, due to their multi-component nature and systemic actions. Studies conducted on *Triphala* and *Nishamalaki* have shown the importance of network pharmacology approaches for studying *Ayurveda* formulations [41]. NK being a formulation used for multiple diseases conditions, we hypothesised a possible network pharmacology mode of action. The analysis of the phytoactives identified from *in silico* data mining, unveiled 1555 proteins as the possible targets. The hub proteins with maximal interaction included carbonic anhydrases (like CA2, CA7, CA4, CA12 and CA12) and proteins like epidermal growth factor receptor (EGFR), aldose reductase (AKR1B1), Matrix metalloproteinases (MMPs) and Aryl hydrocarbon receptor (AR). All these proteins are shown to have significant involvement in diabetic complications. The carbonic anhydrases with higher degree values (CAs) are a group of ubiquitously expressed metalloenzymes, which are involved in numerous physiological processes like gluconeogenesis, lipogenesis, ureagenesis and tumorigenicity [42] [43]. Some of the recent studies indicate their potential as novel targets in obesity and diabetes management [44]. In addition to the above proteins, DPP4, is also identified as a hub protein having multiple protein interactions. This demonstrated that NK interactome may possibly involve many targets which are central to diabetic complications resulting in the multi-modal actions of the formulation.

To delve deep into the diabetic specific PPI network of NK, we utilized the EnrichR tool for disease protein association analysis. 1555 protein targets of NK were submitted in the EnrichR tool, from which 614 diabetes specific proteins were extracted using the key words ‘diabetes mellitus’, ‘insulin resistance’ and ‘hyperglycemia’. Since NK is a formulation used for diabetes and related complications, we wanted to see the overlap of NK diabetic network with related conditions like obesity, inflammation, fatty liver, metabolic dysfunction and other diabetic complications. The Venn diagram analysis of the data showed a cluster of 37 common proteins. This included important signalling kinases (AKT1, PIK3Cα and PIK3Cγ), key obesity associated transcription factors (PPARα and PPARβ) and the pro-inflammatory mediators (IL1α, IL10, TNFα, CCL2, TGFβ1, and NFKβ1). This demonstrated the complex interplay between insulin signalling pathways and lipid synthesis which are deregulated in lifestyle diseases like diabetes, obesity and NAFLD. Apart from this, 87 proteins were found to be overlapping between diabetic complications and obesity, which affirms the well-established role of obesity as a predisposing factor for diabetes. Overlapping presence of DPP4 and GLP1R between fatty liver, obesity and diabetic comorbidities stresses the importance of the gut and its incretin axis in the possible polypharmacological drug network of NK.

A Venn analysis was also conducted between the diabetic complications such as retinopathy, neuropathy, nephropathy and myopathy, wound healing and micro and macrovascular complications. From this, we observed that 23 proteins were common among all the diabetic complications. Among these, inflammatory mediators like TNFα, IL10, CCL2, TLR4 and TGFβ1, MMP9 and MMP2 were present, which are associated with the development of insulin resistance and diabetic nephropathy progression respectively [45] [46]. AKR1B1, present in the common cluster of proteins is involved in the conversion of excess glucose into sorbitol and recent studies prove that overexpression of AKR1B1 is involved many diabetic complications like cardiovascular diseases and retinopathy [47]. Oxidative stress induced by chronic hyperglycemia is one of the main pathways for development of macrovascular and microvascular complications [48]. One of the main mechanisms mediated by polyherbal formulations in alleviating diabetes is by reducing oxidative stress by multiple mechanisms [49]. The Venn diagram between the diabetic complications showed the presence of oxidative stress markers like GSTM1, SOD1 and SOD2 in the central cluster, indicating the possibility of NK acting through oxidative stress signalling, Present *in vitro* and *in silico* studies revealed the DPP4 inhibitory activity of NK as well as its potential to modulate a large diabetic interactome. Out of 37 common proteins identified from the Venn analysis, 28 proteins are found to be associated with five well-studied and critical biochemical pathways in the pathogenesis of diabetes, related complications and comorbidities. Among those, the PI3K/AKT signaling pathway regulates cell proliferation, differentiation, metabolism, and cytoskeletal reorganization (Fig-8). Insulin is one of the main ligands for activation of PI3K signaling, through which it governs a variety of physiological processes in adipose, muscle, brain, liver and pancreas. Multiple proteins like AKT1, AKT2, PIK3Cγ, PIK3Cα and PTEN are associated with these two pathways. They are also involved in MAPK signaling, HIF1α signaling, and AGE signaling all of which play important roles in the progression of Fatty liver disease, obesity and diabetic macrovascular and microvascular complications. Cardiovascular complications like cardiomyopathy are caused by lipid accumulation and atherosclerosis, which is also often seen in metabolic disorders. Lipid and atherosclerosis pathway also includes many proteins like TLR4, IL1α, NFKβ, CCL2 and TNFα, all of which are studied for their inflammatory action in diabetes associated disorders. TLR4 and CCL2 are also emerging targets as they play a key role in many diabetic complications like nephropathy and retinopathy. Other important signaling pathways associated with the disease overlap of proteins include HIF1α signaling, cAMP signaling, RAP1 signaling, Calcium signaling and apoptotic pathway.

**Figure-8:**
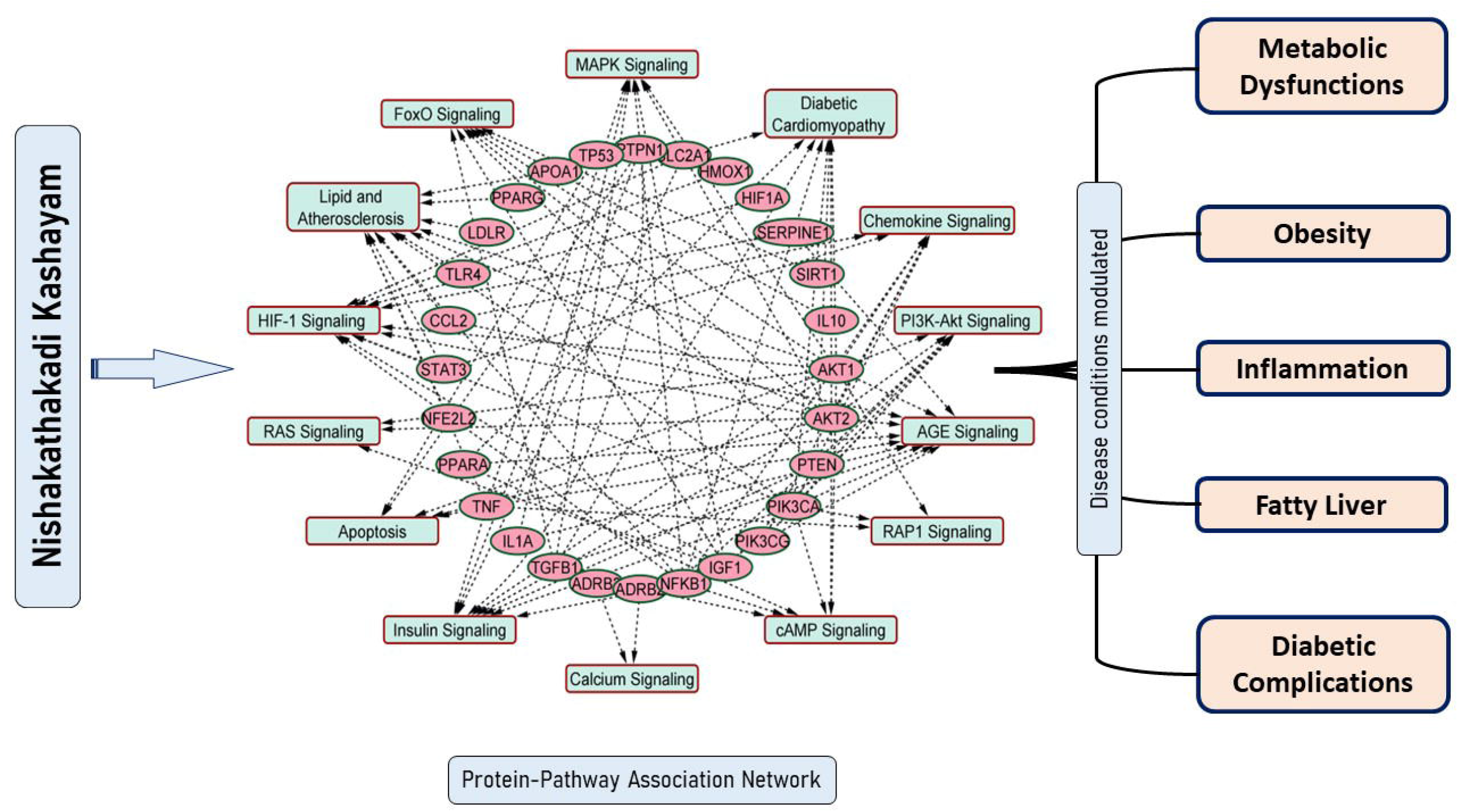
A pictorial depiction of the common protein cluster that emerged from the Venn analysis between the five disease categories potentially modulated by NK and their associated signaling pathways.

## 5. Conclusion

The present work as evidenced through *in vitro* and *in silico* observations suggests that the anti-diabetic formulation NK has a strong DPP4 inhibitory potential. Further, our network pharmacology studies of the phytochemicals revealed a complex molecular interaction network consisting of crucial proteins and significant pathways in diabetes, diabetes-related complications and associated metabolic diseases. This integrated approach of combining *in vitro* and *in silico* methods offers the much-needed scientific rationale for traditionally prescribed polyherbal *Ayurveda* formulations in the treatment of complex diseases like type 2 diabetes. The pathways and molecular targets obtained from our work merits detailed investigation which will enable us to have more insight into the action of this *Ayurveda* formulation.

## Acknowledgement

The authors acknowledge the financial support received by Anjana from Tata Education and Development Trust and Rural India Support Trust (RIST). Authors also acknowledge the financial support for a Research Fellow from TDU, Bangalore. The authors also acknowledge the DS Endowment support received from TDU for the project. The authors acknowledge SASTRA Deemed to be University for Schrodinger software (https://www.schrodinger.com/) support. The authors Sthitaprajna Sahoo and Abhijnan Chakraborty contributed to the work as part of their MentX virtual internship program under the guidance of Suma Mohan S.

## Abbreviations

AKR1B1: Aldo-keto reductase family 1 member B
AKT1: AKT serine/threonine kinase 1
AKT2: AKT serine/threonine kinase 2
CA2: Carbonic anhydrase 1
CA7: Carbonic anhydrase 2
CA4: Carbonic anhydrase 4
CA12: Carbonic anhydrase 12
CCL2: C-C motif chemokine ligand 2
DPP4: Dipeptidyl peptidase 4
EGFR: Epidermal growth factor receptor
GLP1R: Glucagon like peptide 1 receptor
GSTM1: glutathione S-transferase mu 1
IL1α: Interleukin 1 alpha
IL10: Interleukin 10
MMP9: Matrix metallopeptidase 9
MMP2: Matrix metallopeptidase 2
PPARα: Peroxisome proliferator activated receptor alpha
PPARβ: Peroxisome proliferator activated receptor beta
NFKβ1: Nuclear factor kappa B subunit 1
PIK3Cα: Phosphatidylinositol-4,5-bisphosphate 3-kinase catalytic subunit alpha
PIK3Cγ: Phosphatidylinositol 4,5-bisphosphate 3-kinase catalytic subunit gamma isoform
PTEN: Phosphatase and tensin homolog
SOD1: Superoxide dismutase 1
SOD2: Superoxide dismutase 2
TGFβ1: Transforming growth factor beta 1
TLR4: Toll like receptor 4

